# Label free sensing with Terahertz multiple ring resonators

**DOI:** 10.1101/2021.03.07.434261

**Authors:** Xaing Bai, Lujun Zhe

**Affiliations:** School of Electrical and Computer Engineering, Xiangtan University

**Keywords:** Terahertz sensing, Graphene, Surface plasmon polaritons, Metamaterials, Ring resonators, Terahertz spectroscopy

## Abstract

Apart from their relevance for spectroscopy and imaging, terahertz signals have attracted a lot of interest for sensing. In this paper, a label free terahertz sensor is proposed, which can be employed to detect the presence of molecules in the environment. The proposed sensor consists of an array of ring resonators, resonating at the frequency of *f=1*.*2 THz*. By providing full-wave numerical simulations, it is shown that the proposed sensor is able to sense the variation of the refractive index in the environment. The proposed structure is found to show larger sensitivity compared to previous reports. Our findings provide a novel platform to realize label free terahertz sensors with extremely large sensitivity.

## I. Introduction

THz band radiation is a part of the electromagnetic spectrum, occupying the frequency between *f* = 0.1 − 10 *THZ* [1-10]. The waves in this frequency have some common properties. For instance, they can penetrate into various kinds of dielectric materials. Due to the lack of efficient sources and detectors, this frequency spectrum is called the THz gap. Remarkably, the energy of photons in the THz is smaller than the bandgap of the dielectric materials. Therefore, THz waves can penetrate in these materials. The transmitted wave can be employed to characterize the properties of materials, allowing one nondestructive evaluation, security and biomedical sensing [11-20].

One important application of terahertz waves is label-free sensing [], which can be used for molecular sensing, for instance. The usage of label-free technique prevents the challenges associated with labeling, including preparation of the sample. One of the most important way for label free sensing is based on refractive index changes. In this method, the structure senses biomolecules presence directly from the variation of the refractive index of the cladding medium. Many previous reports exist on the optical structures to achieve label-free sensing. One of the most important classes of such sensors are surface plasmon resonance (SPR) based sensors [21-25].

The working principle of the SPP sensors are the following: since the power of surface plasmon modes is mainly confined in an interface between metal and dielectric, it is sensitive to the changes in the refractive index of cladding material.

The oscillation of the electrons of a material on the edge of an interface made of a dielectric and metal is the basic principle of surface plasmon polaritons at any frequency range. The propagation loss of the surface plasmon polaritons in the spectrum is high. However, their localization is appropriate for sensing the changes in the dielectric refractive index. While various sensors are being created these days, the THz frequency with its exotic features is of utmost interest for biosensor researchers. One important property is that the associated phononic modes of many macro molecules like DNAs are in this particular range. This makes terahertz sensors perfect candidates to detect chemical matters more efficiently [8]. The absorption loss of THz SPPs on the interface of the two media is much smaller compared to the visible surface plasmon polaritons. Yet, they have very small localization to a dielectric material. As a result, surface plasmon polaritons are not appropriate for integration. This issue can be addressed in various approaches. Replacing metals with graphene strips can be a solution to solve this problem.

Graphene [26-36] is a thin layer carbon atom. The atoms of the monolayer are tightly packed into a two-dimensional hexagonal lattice of atoms. In particular, graphene can be considered as the two dimensional version of the bulky graphite. Research has been demonstrated that, in THz frequency regime, the surface conductivity of graphene flakes become imaginary, a property which allows the graphene strip to exhibit TM plasmonic modes. These modes have a much smaller effective wavelength than air, enabling miniaturization. Importantly, the graphene surface conductivity can be tuned dynamically over a wide range of frequencies, by just adding a bias or chemical doping. As a result, the propagation constant corresponding to the plasmonic mode can be dynamically adjusted. This enables reconfiguration of the associated device [37-46]. Moreover, confining electromagnetic fields in a short volume than regular materials has made graphene as a perfect choice for realization of nano structures.

In this paper, it is proposed to perform label free sensing by means of an array of surface plasmon polariton ring resonators. The ring resonators of the array are made of graphene strips sandwiched between two dielectric layers with different refractive indexes. It is shown that the proposed structure is capable of sensing the changes in the refractive index of the environment through a shift in the resonance frequency of the ring resonator.

By providing the results of our finite element time domain simulations, it is shown that the proposed structure show a larger sensitivity compared to previous reports. It is further investigated that by tuning the chemical potentials of the graphene strips, one can reconfigure the proposed sensor to different frequencies. The chemical potentials of the graphene strips can be easily reconfigured using a change in the voltage applied to it. Alternatively, it is possible to change the chemical potentials of the graphene strips using chemical doping. We further investigate the effect of this change in the chemical potentials on the sensitivity and figure of merit of the proposed sensor. The proposed results provides a new route for designing ultra-sensitive label free Terahertz sensing devices, whose sensitivity and other associated parameters may be tuned in a dynamic way.

## II. Proposed structure

The proposed structure is shown in Fig.1. As it is observed, the proposed structure includes an array of split ring resonators. An optical waveguide excites the structure. Each ring resonator consists of a graphene strip, sandwiched between two dielectrics, namely Si and SiO_2_. The entire structure is placed on top of a silicon substrate.

**Fig. 1:**
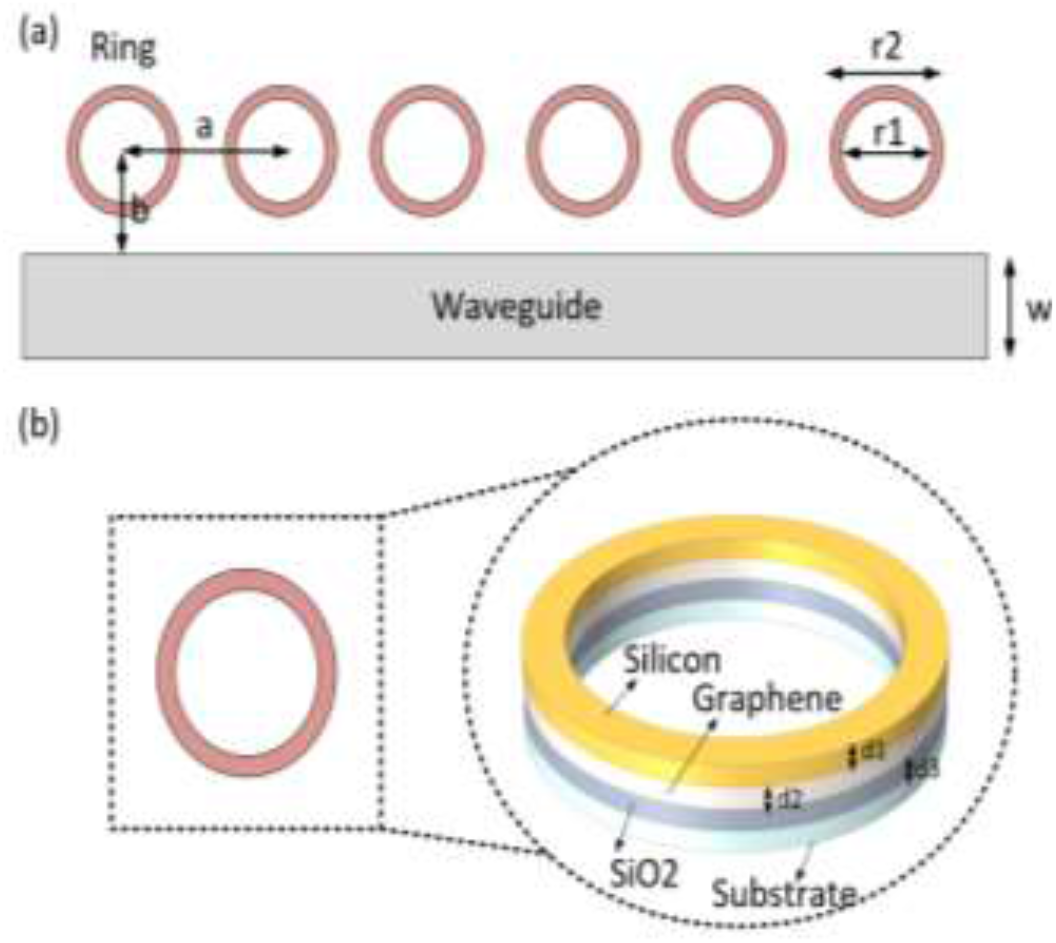
a) Proposed structure for performing THz label free sensing. The structure is composed of an array of ring resonators. B, structure of the ring resonators of the array

In order to characterize the structure, we use Lumerical software, a full-wave electromagnetic solver based on finite domain time difference method (FDTD). The graphene strips in our structure is modeled based on an impedance layer. The surface conductivity of this impedance layer is defined by the conductance of the sheet, determined by the so-called Kubo formula.

The dielectrics of the structure are modeled using a blank material whose refractive index is in accordance with the permittivity of the material. The corresponding transmission coefficient of the structure is numerically calculated and shown in Fig. 2. As observed, the obtained result shows a resonance at the resonance frequency of the structure.

**Fig. 2:**
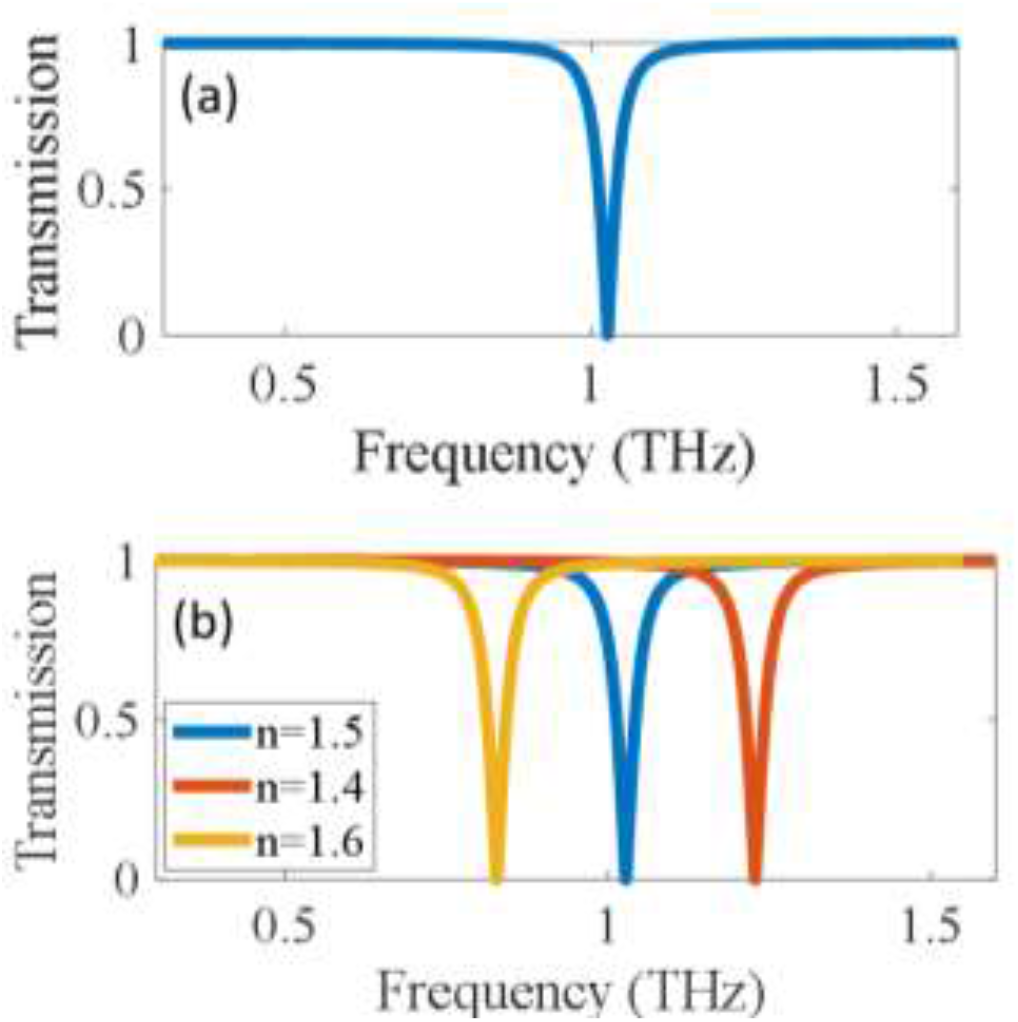
a) Transmission spectrum of the proposed structure (b) variation of the transmission spectrum as a function of the refractive index of the top layer

The obtained resonance is then used to perform label free sensing as it is described in what follows. Suppose that the presence of the molecules in the environment causes a change in the refractive index of the environment. This change can create a shift in the spectrum of the system. This effect is demonstrated in Fig. 3 of the manuscript, where the variation of the transmission spectrum of the system as a function of the refractive index of the cladding, i.e. the environment surrounding the structure is reported. As it is observed, the change in the refractive index has led to a shift in the spectrum. This shift can be used to perform label free sensing. In order to unambiguously investigate the sensitivity of the sensor under study, the variation of the resonance frequency shift as a function of the change in the refractive index is calculated and reported in Fig. 4. The figure reports this variation for various values of chemical potentials. The slope of these curves determines the sensitivity of sensor for that particular chemical potential.

**Fig. 3:**
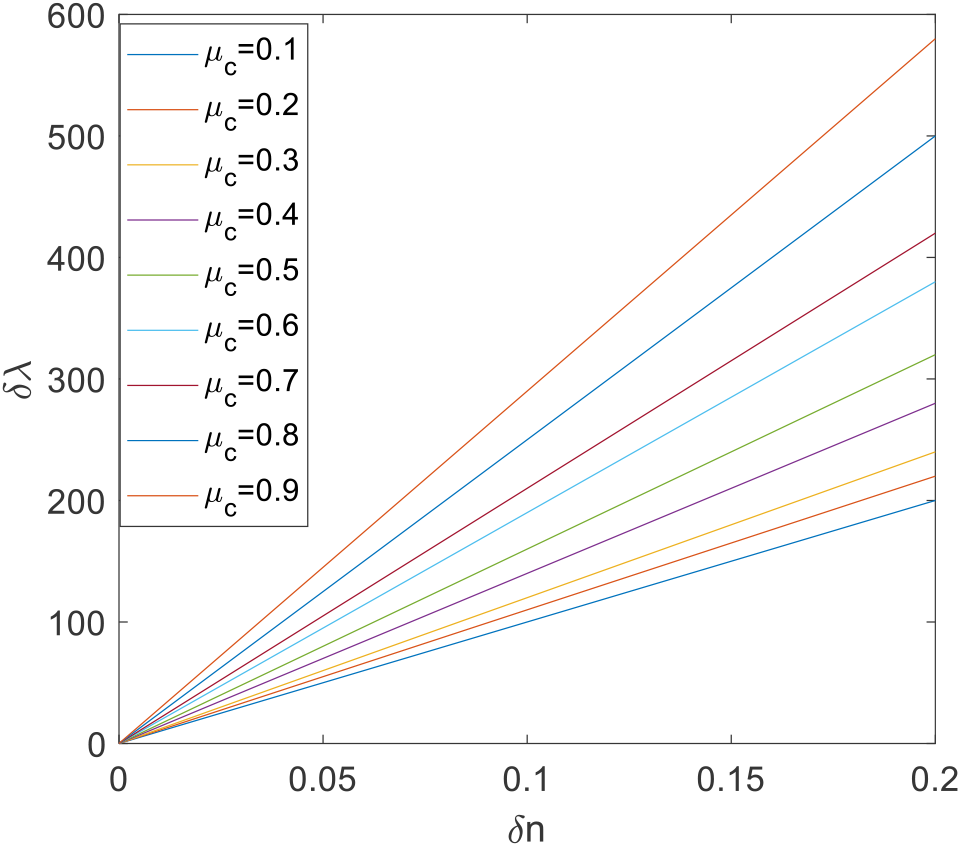
Sensitivity of the proposed sensor as a function of the Fermi level of graphene layers. The sensitivity is determined by the slope of the curves

To assess the functionality of the sensor, the quality factor of the sensor is another important factor that must be considered. The parameter Q can be obtained making use of 3dB bandwidth. As it is known, one common technique to obtain the quality factor of a resonator is based on using the complex resonance frequency. The quality factor of the obtained resonators were achieved to be *Q= 1895*.

Note that, in general, there exist a tradeoff between the sensitivity and quality factor of the proposed sensor. In particular, one requires to determine if the high Q is the important factor or a higher sensitivity matters. The Figure of Merit (FOM) is a parameter that can be considered as a perfect factor which takes into account both of these parameters. The FOM of the sensor under study is calculated to be FOM=96. To conclude, in this paper, we discussed designing a label free THz sensor. The proposed sensor was based on an array of graphene ring. By providing the results of our full-wave numerical simulations, we showed that the proposed sensor is capable of sensing the changes in environment through a shift in the spectrum of the system. The designed system is a appealing example of a terahertz bio sensor that provides a large sensitivity, quality factor as well as figure of merit. The generalization of the approach to the case of ring resonators supporting Fano or Kerkerr resonances can be a trivial extension of this research work [47-49].

